# Infection of human lymphomononuclear cells by SARS-CoV-2

**DOI:** 10.1101/2020.07.28.225912

**Authors:** Marjorie C Pontelli, Italo A Castro, Ronaldo B Martins, Flávio P Veras, Leonardo La Serra, Daniele C Nascimento, Ricardo S Cardoso, Roberta Rosales, Thais M Lima, Juliano P Souza, Diego B Caetité, Mikhael H F de Lima, Juliana T Kawahisa, Marcela C Giannini, Letícia P Bonjorno, Maria I F Lopes, Sabrina S Batah, Li Siyuan, Rodrigo L Assad, Sergio C L Almeida, Fabiola R Oliveira, Maíra N Benatti, Lorena L F Pontes, Rodrigo C Santana, Fernando C Vilar, Maria A Martins, Thiago M Cunha, Rodrigo T Calado, José C Alves-Filho, Dario S Zamboni, Alexandre Fabro, Paulo Louzada-Junior, Rene D R Oliveira, Fernando Q Cunha, Eurico Arruda

**Author notes:** Corresponding authors: Marjorie Cornejo Pontelli; Virology Research Center, Ribeirao Preto Medical School, 3900 Bandeirantes Av, 14049-900, Ribeirao Preto –SP – Brazil., Italo Araujo Castro; Virology Research Center, Ribeirao Preto Medical School, 3900 Bandeirantes Av, 14049-900, Ribeirao Preto –SP – Brazil., Eurico Arruda; Virology Research Center, Ribeirao Preto Medical School, 3900 Bandeirantes Av, 14049-900, Ribeirao Preto –SP – Brazil. Contributed equally to the study (Marjorie C Pontelli, Italo A Castro).

## Abstract

Although SARS-CoV-2 severe infection is associated with a hyperinflammatory state, lymphopenia is an immunological hallmark, and correlates with poor prognosis in COVID-19. However, it remains unknown if circulating human lymphocytes and monocytes are susceptible to SARS-CoV-2 infection. In this study, SARS-CoV-2 infection of human peripheral blood mononuclear cells (PBMCs) was investigated both *in vitro* and *in vivo*. We found that *in vitro* infection of whole PBMCs from healthy donors was productive of virus progeny. Results revealed that monocytes, as well as B and T lymphocytes, are susceptible to SARS-CoV-2 active infection and viral replication was indicated by detection of double-stranded RNA. Moreover, flow cytometry and immunofluorescence analysis revealed that SARS-CoV-2 was frequently detected in monocytes and B lymphocytes from COVID-19 patients, and less frequently in CD4^+^T lymphocytes. The rates of SARS-CoV-2-infected monocytes in PBMCs from COVID-19 patients increased over time from symptom onset. Additionally, SARS-CoV-2-positive monocytes and B and CD4+T lymphocytes were detected by immunohistochemistry in post mortem lung tissue. SARS-CoV-2 infection of blood circulating leukocytes in COVID-19 patients may have important implications for disease pathogenesis, immune dysfunction, and virus spread within the host.

## Introduction

In December 2019, a new coronavirus emerged as the cause of a severe acute respiratory disease named Coronavirus-related disease 2019 (COVID-19). The virus that spilled over to humans in China was classified in the family *Coronaviridae*, genus *Betacoronavirus*, and was named Severe Acute Respiratory Syndrome Coronavirus 2 (SARS-CoV-2), for its similarity to SARS-CoV [1].

Since its emergence, SARS-CoV-2 has spread to 185 countries/political regions and infected more than 11 million people worldwide, with a death toll of approximately 500,000 cases. The main clinical features of COVID-19 are fever, dry cough, dyspnea and myalgia, but some patients rapidly evolve to severe respiratory distress syndrome [2].

Previous studies have shown that inflammatory cytokine storm and lymphocytopenia are important markers of severe COVID-19 cases, with severe functional exhaustion of TCD4+ and TCD8+ lymphocytes [2–4]. Interestingly, peripheral blood mononuclear cells (PBMCs) from COVID-19 patients showed upregulation of autophagy and apoptosis pathways [5], suggesting that dampening of the immune system by SARS-CoV-2 infection may have a strong impact on the clinical outcome of severe COVID-19.

A decrease in circulating lymphocytes has been associated with poor COVID-19 outcome, but it is still unclear whether that lymphopenia is directly due to SARS-CoV-2 infection of lymphocytes with consequent cell death. SARS-CoV-2 interacts with target cells via binding of its major surface glycoprotein spike (S) with the angiotensin-converting enzyme 2 (ACE2) present in the cell membrane [6]. ACE2-independent cell-entry has also been reported and could be an alternative mechanism of SARS-CoV-2 entry in cells with low ACE2 expression [7]. Cleavage of the S protein is required for efficient entry of SARS-CoV-2, which is accomplished by transmembrane serine protease TMPRSS2 [6].

In addition to respiratory disease, COVID-19 patients frequently develop gastrointestinal symptoms, which is in keeping with the high expression of TMPRSS2 and ACE2 documented in enterocytes [8]. Furthermore, the SARS-CoV-2 antigen was found *post mortem* in the spleen and lymph nodes with pathological signs of damage. In these organs, monocytes do contain viral antigens, but it was not clear whether this was due to active viral replication or phagocytosis, nor if monocytes become infected before reaching secondary lymphoid tissues [9]. While SARS-CoV-2 causes viremia, until now, infectious SARS-CoV-2 was not successfully isolated from peripheral blood in COVID-19 patients, and it is suggested that the virus in blood may be cell-associated [5, 9]. In this study, we investigated the susceptibility and permissiveness of human peripheral blood mononuclear cells (PBMC) to SARS-CoV-2. We found that PBMCs are susceptible and permissive to SARS-CoV-2 infection, both *in vivo* and *ex vivo*, which seems to play a direct role in the reduction of circulating lymphocytes.

## Patients and Methods

### Ethical statement and COVID-19 patients

The study was approved by the National Ethics Committee (CONEP, CAAE: 30248420.9.0000.5440 and 31797820.8.0000.5440). A total of 29 hospitalized patients were enrolled, all with clinical and radiological features of COVID-19 and confirmed SARS-CoV-2 infection by RT-PCR in respiratory secretions, with detection of specific IgM or IgG antibodies to SARS-CoV-2. Clinical features, laboratory results and drug therapies are summarized in Supplementary Table 1. For all comparisons, 12 age and gender-matching healty controls were also enrolled. Written informed consent was obtained for both patients and healthy controls.

### Production of mouse anti-SARS-CoV-2 hyperimmune serum

Male C57Bl/6 mice were bred and maintained under specific pathogen-free conditions at the animal facility of the Ribeirão Preto Medical School (FMRP) at University of São Paulo. The protocol for production of mouse hyperimmune serum were carried out with 8-week-old male mice following the institutional guidelines on ethics in animal experiments and was approved by the University of São Paulo Ethics Committee for Animal Experimental Research -CETEA (Protocol no. 001/2020-1). To immunize animals, virus stock was inactivated by adding formaldehyde to a final concentration of 0.2%, and incubated overnight at 37°C. Then, virus was purified by ultracentrifugation (10% sucrose cushion, 159.000 × g for 1h). The pellet was resuspended with Phosphate Buffer Saline (PBS) 1x and stored at -20°C. In order to confirm inactivation, titration of the inactivated product was done both by TCID_50_ and by plaque assay in Vero-E6 cells with 5-day incubation, without any cytopathic effects. Three C57Bl/6 mice were inoculated intramuscularly with an emulsion containing the equivalent of 10^6^ TCID_50_ of inactivated SARS-CoV-2 in complete Freund’s adjuvant (CFA, BD, cat. 263810) diluted 1:1 in PBS. Boosts were given with inactivated SARS-CoV-2 in incomplete Freund’s adjuvant (without *M. tuberculosis,* IFA, BD, cat. 263910) on days 7 and 14 after the first immunization. One week after the last dose, animals were euthanized with an excess of anesthetics xylazine (60 mg/kg) and ketamine (300 mg/kg), following exsanguination by cardiac puncture. Animal serum conversion was evaluated by indirect immunofluorescence using slide preparations of SARS-CoV-2 infected Caco-2 cells, fixed with 4% paraformaldehyde and AlexaFluor 488-labelled rabbit anti-mouse secondary antibody. Coverslips were analyzed using an optic microscope (Olympus BX40).

### Isolation of peripheral blood mononuclear cells (PBMCs)

Human PBMCs were isolated from COVID-19 patients or healthy donors by density gradient using Percoll (GE Healthcare, cat. 17-5445-01), as previously described [10, 11]. PBMCs were washed, resuspended in RPMI 1640 supplemented with 10% fetal bovine serum (FBS) and kept on ice until further use.

### Virus and cell lines

The passage 1 (P1) of SARS-CoV-2 Brazil/SPBR-02/2020 isolate obtained in Vero-E6 from a COVID-19 patient in Sao Paulo was kindly provided by Prof. Edison Durigon (ICB-USP). P1 was diluted 1:1000 in Dulbecco’s modified Eagle’s medium (DMEM) and inoculated in Vero-E6 cells monolayers to produce the P2 stock. For stock titration, serial 10-fold dilutions were inoculated in quadruplicate monolayers of Vero-E6 cells and incubated at 37°C in 5% CO_2_. On the fourth day of incubation, the presence of cytopathic effect (CPE) was recorded **(Supplementary Fig 1A)** and titers were expressed as the 50% tissue culture infectious dose (TCID_50_), using the Reed-Muench method. All experiments involving SARS-CoV-2 propagation were done in biosafety level 3 laboratory.

**Figure 1.**
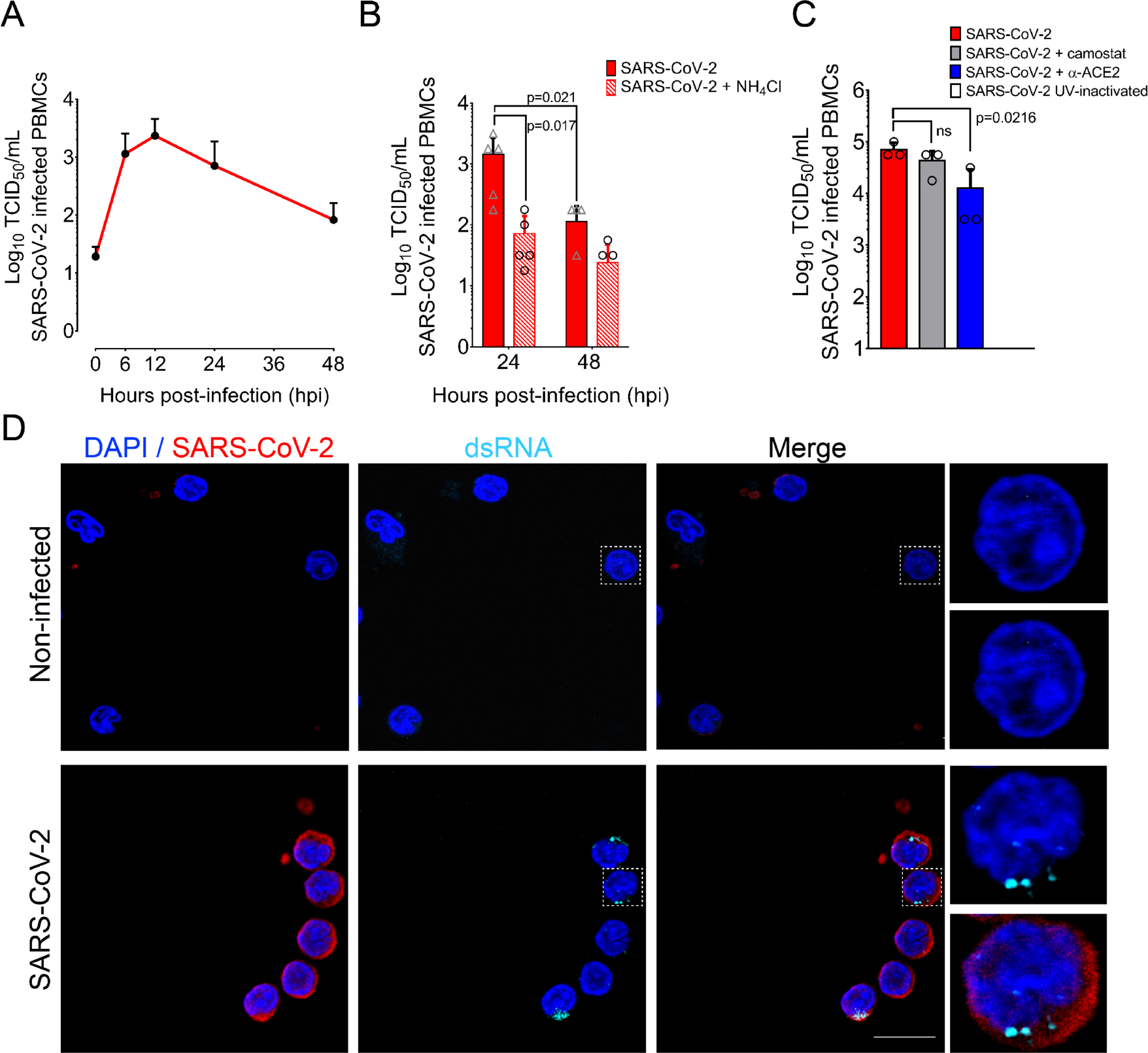
Human primary blood cells are susceptible and permissive to SARS-CoV-2. Blood from five healthy donors was collected and PBMCs were separated by Ficoll density gradient. Cells were infected with SARS-CoV-2 Brazil/SPBR-02/2020 (MOI-1) and cultured for 48 h. (A) Overtime virus progeny production from PBMCs infected with SARS-CoV-2. Supernatants from cultured PBMCs were collected at each time point and titrated by TCID_50_. The small symbols represent individual values (5 healthy donors) and error bars depict standard deviation. (B) SARS-CoV-2 progeny titers in supernatants of infected PBMCs at 24 and 48 hpi, with and without treatment with 20mM NH_4_Cl. (C) Effects of blocking SARS-CoV-2 cell receptor ACE2 and TMPRSS2 on virus progeny production. Infected PBMCs were exposed to antibody anti-ACE2 or Camostat and virus progeny was titrated in supernatants at 24 hpi. (D) Immunostaining for dsRNA in PBMCs cultured on poly-lysine –coated coverslips 6h after SARS-CoV-2 infection. Cells were fixed, immunostained for SARS-CoV-2 (red), dsRNA (cyan) and analyzed by confocal microscopy. Statistical analysis was performed using one-way or two-way ANOVA. Tukey’s or Holm-Sidak post-tests were applied when suitable. P values < 0.05 were considered significant. Magnification: 63x. Scale bars 10 µM.

### *In vitro* infection of PBMCs

For these experiments, 10^6^ PBMCs from 5 healthy donors were infected with SARS-CoV-2 (MOI=1) in RPMI with 0% FBS at RT for 1 h under orbital agitation. Next, cells were pelleted at 300 × g, the inoculum was washed and replaced by RPMI with 2% FBS and cells were incubated at 37°C in 5% CO_2_. As controls, equivalent quantities of cells were exposed to UV-inactivated SARS-CoV-2 and treated in the same way. We also used a control consisting of cells treated with 20 mM of NH_4_Cl starting 20 min before infection and maintained throughout the entire incubation period. Supernatants from PBMCs were collected at 0, 6, 12, 24 and 48 h post-infection and subjected to serial ten-fold dilutions to determine virus titers by TCID_50_, as previously described. In parallel, PBMCs were treated with 0.5ug/ml Camostat (Sigma Aldrich, cat. SML005) or with 10 uM of anti-ACE2 antibody (Rhea Biotech, cat. IM-0060) starting 1 hour before infection. Then, innoculum was washed, the media containing the different treatments were replaced, and the PBMCs were kept for 24h at 37°C in 5% CO_2_.

### RNA extraction and real-time RT-PCR

SARS-CoV-2 RNA detection was done with primer-probe sets for SARS-CoV-2 according to the USA-CDC protocol, targeting the virus N1 gene, and using the RNAse-P housekeeping gene as control, by one-step real-time RT-PCR. Total RNA was extracted with Trizol® (Invitrogen, CA, EUA) from 250µL of homogenized cell pellets and supernatants from in vitro assays. All real-time PCR assays were done on a Step-One Plus thermocycler (Applied Biosystems, Foster City, CA, USA). Briefly, after Trizol® extraction, 100 ng of RNA was used for genome amplification with N1 primers (20 µM) and probe (5 µM), and TaqPath 1-Step qRT-PCR Master Mix (Applied Biosystems, Foster City, CA, USA), with the following parameters: 25°C for 2 min, 50°C for 15 min, 95°C for 2 min, followed by 45 cycles of 94 °C for 5 s and 60°C for 30s. Viral loads of SARS-COV-2 were determined using a standard curve prepared with a plasmid containing a 944bp amplicon, which includes all three targets for the sets of primers/probes designed by CDC protocol (N1, N2 and N3), inserted into a TA cloning vector (PTZ57R/T CloneJetTM Cloning Kit Thermo Fisher^®^). Results of viral RNA quantifications by one-step qRT-PCR were plotted with GraphPad® Prism 8.4.2 software.

### Indirect immunofluorescence staining of SARS-CoV-2 infected cells

Coverslips pre-treated with poly-lysine 0.1% (Sigma-Aldrich, cat. P8920) were incubated with isolated PBMCs from patients or healthy donors at 37°C, 20 minutes for cell adherence. After that, coverslip-containing cells were fixed with 4% paraformaldehyde (PFA) in PBS for 15 minutes, and then washed 3 times with PBS. To detect viral antigens in cells, we used serum from a recovered COVID-19 patient, which was first tested for specificity by immunofluorescence in SARS-CoV-2 infected Vero CCL81 cells (**Supplementary Fig 1B).** In addition, for each experiment using the referred serum we included cells from healthy donors or non-infected cells. As an isotype control of this serum, we used a human serum collected in 2016. Biotin-conjugated anti-human IgG (Sigma-Aldrich, cat. B-1140) was used as the secondary antibody, followed by amplification with the TSA Cyanine 3 System (Perkin Elmer, NEL704A001KT), following the manufacturer’s protocol. To determine the phenotype of SARS-CoV-2-infected cells, we used primary antibodies for CD4 (Abcam cat. ab133616), CD8 (Abcam cat. ab4055), CD14 (Abcam cat. ab133335), CD19 (Abcam cat. ab134114), CD20 (Abcam cat. ab103573). For detection of virus replication, we used a mouse anti-dsRNA J2 (dsRNA; English & Scientific Consulting Kft, Hungary), which binds to dsRNA of 40 bp or longer. Secondary antibodies used were polyclonal anti-rabbit conjugated with 488 (Thermo Fisher cat. A21202), 594 (Abcam cat. ab150116) or 647 (Abcam cat. ab150079). The Golgi complex and nuclei staining were carried out using a mouse anti-GM130 (BD cat. 610822) and 4′,6-diamidino-2-phenylindole dihydrochloride dye (DAPI, Thermo Fisher cat. 62248), respectively.

### Confocal microscopy

PBMCs from confirmed COVID-19 patients and from healthy control donors were stained with human serum containing antibodies to SARS-CoV-2, and with commercial antibodies to the different cell phenotypes, followed by the appropriate secondary antibodies. Preparations were analyzed in a Zeiss Confocal 780 microscope in a Tile 3×3 in a single focal plane. The quantity of SARS-CoV-2-positive cells of different phenotypes was quantified by using the analyze particles tool from Fiji by ImageJ.

### Flow cytometry

Unseparated whole blood leukocyte samples from COVID-19 patients or healthy donors infected in vitro with SARS-CoV-2 were surface stained with Fixable Viability Dye eFluor™ 780 (eBioscience) and monoclonal antibodies specific for CD3 (APC eBioscience cat. 17-0036-42), CD4 (PerCP-Cy5.5 BD cat. 560650), CD8 (PE-Cy7 BD cat. 557746), CD19 (APC BioLegend cat. 302212), CD14 (PerCP Abcam cat. ab91146), CD16 (PE eBioscience cat. 12-0168-42), CCR2 (BV BioLegend cat. 357210) for 30 min at 4°C, according to manufacturer’s instructions. Detection of SARS-CoV-2 by flow cytometry was performed with BD Cytofix/Cytoperm™ kit to enable access to intracellular antigens using mouse polyclonal antibody raised against formalin-inactivated SARS-CoV-2, as described early in this manuscript, for 15 min at 4°C. To ensure that viral detection was specific for replicating intracellular viruses, additional preparations of infected cells were stained without permeabilization. Treatment with trypsin for 60 min on ice after infection to remove surface-bound viral particles was also included as a second control (**Supplementary Fig 2)**. SARS-CoV-2 antibodies were detected with secondary anti-Mouse Alexa488. Surface phosphatidylserine (PS) staining was carried out in whole blood using ApoScreen AnnexinV-FITC apoptosis kit (SouthernBiotech cat 10010-02), following manufacturer’s guidelines. All data were acquired using a Verse or Canto flow cytometers (BD Biosciences) and subsequent analysis was done using FlowJo (TreeStar) software. Gating strategies are illustrated in **Supplementary Fig 3**.

**Figure 2.**
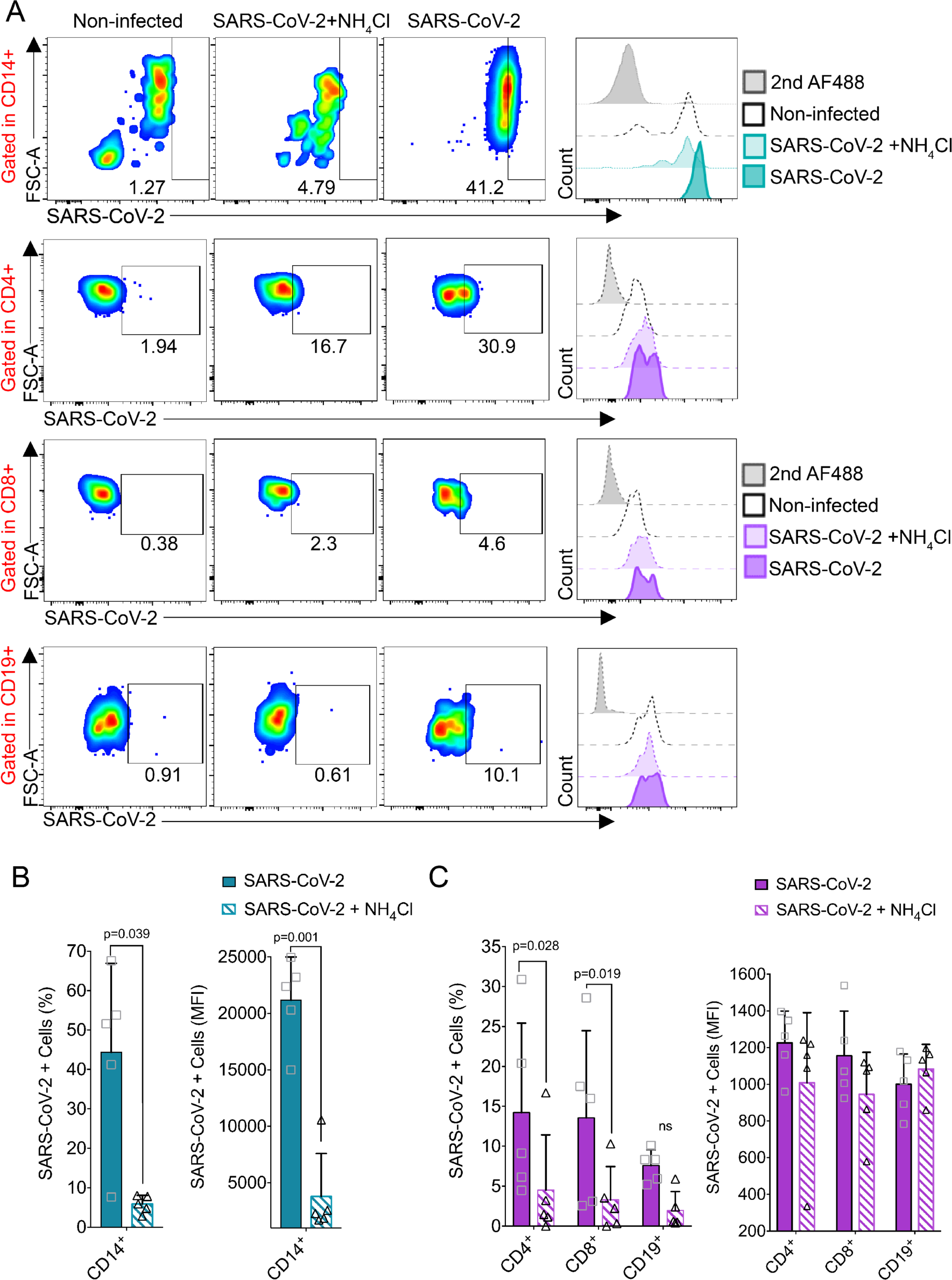
SARS-CoV-2 differentially infects subsets of human PBMCs in vitro. (A) Representative flow cytometry plots of PBMCs infected with SARS-CoV-2 (24 hpi) in the presence or absence of 20mM NH_4_Cl with gating in live CD14^+^, CD4^+^, CD8^+^ or CD19^+^ cells. Representative histograms of the fluorescence for each condition in comparison with the proper controls. The gray-shaded curve indicates secondary antibody Alexa488 signal background, while the dashed curve indicates the background signal in mock-infected cells. The light- and dark-colored curves indicate respectively cells infected in the presence and absence of NH_4_Cl. Percentages of SARS-CoV-2-infected monocytes (B) and lymphocytes (C), showing the average mean fluorescent intensity (MFI) in the panels on the right and the frequency (%) of SARS-CoV-2-infected cells in the panels on the left. Mean ± s.d. is indicated on the bar graphs. Significance was determined by one-way ANOVA with Bonferroni’s post-test.

**Figure 3.**
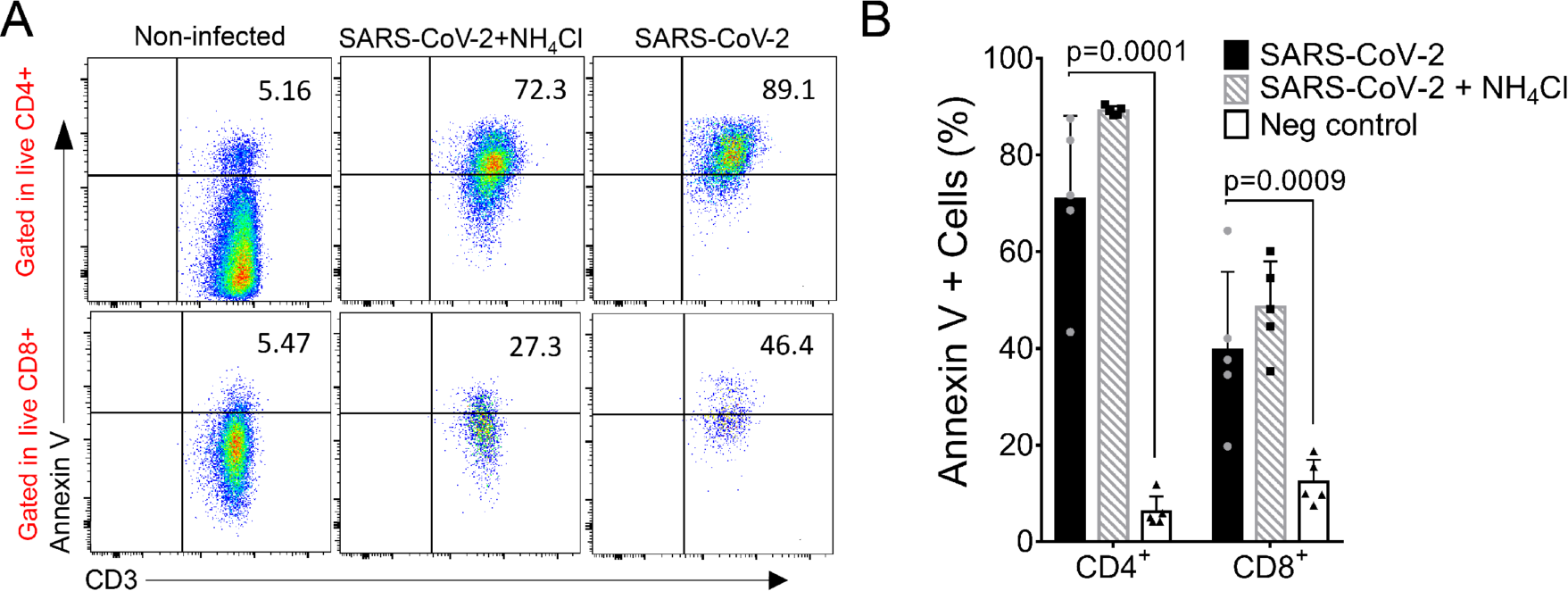
SARS-CoV-2 infection of PBMCs increases expression of phosphatidylserine (PS) in T lymphocytes. (A) Representative flow cytometry plots of live CD4^+^ and CD8^+^ T cells positive for Annexin V staining in PBMCs from five heathy donors 24h after infection with SARS-CoV-2 (MOI= 1). (B) Percentages of live lymphocytes positive for Annexin V and expressing PS in the cell surface after SARS-CoV-2 *in vitro* infection. Mean ± s.d. is indicated for all bar graphs. Significance was determined by one-way ANOVA and Bonferroni’s post-test.

### Serial immunohistochemistry

Tissue sections from paraffin-embedded lung fragments obtained from two COVID-19 fatal cases were tested by immunohistochemistry (IHC) using anti-SARS-CoV-2 polyclonal antibody for in situ detection of SARS-CoV-2. Sequential immunoperoxidase labeling and erasing (SIMPLE) [12] was then performed to determine the immunophenotypes of SARS-CoV-2 infected cells, using antibodies to CD4 (Abcam cat. ab133616), CD20 (Abcam cat. ab103573), CD14 (Abcam cat. ab133335) and IL-6 (BD cat. 554400). After each round of staining, slides were scanned using a VS120 ScanScope (Olympus) under 400x magnification. Images were pseudocolored and overlaid in the first image of the preparation counterstained with hematoxylin using ImageJ v1.50b (NIH, USA) and Adobe Photoshop CS5 software (Adobe Systems, San Jose, CA, USA). Lung paraffin-embedded tissue obtained from a fatal case of hantavirus infection in 2016 was used as a negative control for SARS-CoV-2 staining.

### Statistical analysis

All descriptive statistics, patient stratification, and positive cell frequencies were done using GraphPad Prism Software, version 6.0. Correlation analysis, one-way ANOVA, two-way ANOVA, linear regressions, Holm-Sidak, and Bonferroni post-tests were also performed using GraphPad Prism. Values of *P* <0.05 were considered significant, as described in all figures.

## Results

### SARS-CoV-2 infection of human PBMCs is productive

Considering that human lymphocyte and monocyte lineages are susceptible to SARS-CoV-2 infection *in vitro*, we sought to determine whether primary cultures of human PBMCs could also be infected. Therefore, PBMCs from five healthy donors were infected *in vitro* at a MOI=1. After 0, 6, 12, 24 and 48 hpi, supernatants were harvested, and virus progeny was titrated. SARS-CoV-2 titers peaked between 6 and 12 hpi, resulting in a 100-fold increase from the initial input, and decreased steadily thereof (**Fig 1A**). As expected, induction of general intracellular alkalization by treatment with NH_4_Cl reduced progeny production by approximately 10x (p=0.017). Interestingly, virus progeny production was not entirely abolished by NH_4_Cl treatment, suggesting an entry pathway alternative to endosomal acidification in PBMCs (**Fig 1B**).

Even though expression of ACE2 is minimal in human PBMCs in general [13, 14], we evaluated the viral production after blocking ACE2 and TMPRSS2. Virus titers obtained after Camostat blockage of TMPRSS2 were not significantly different from those obtained without the treatment (**Fig 1C**), suggesting that PBMC infection is not dependent on TMPRSS2. Conversely, the blockage of ACE2 with anti-ACE2 antibody resulted in reduction, but not abrogation of SARS-CoV-2 progeny production after 24 hpi (p=0.0216) (**Fig 1C**), indicating that SARS-CoV-2 can infect human PBMCs independently of ACE2.

Coronavirus replication entails the formation of abundant double-stranded RNAs (dsRNA) in the cytoplasm of infected cells, and thus its intracellular detection is a reliable marker of viral replication. Therefore, infected PBMCs were stained for SARS-CoV-2 and dsRNA and analyzed by confocal microscopy. Most SARS-CoV-2-positive cells were also positive for dsRNA, and rates of double-positive cells counted at 6 hours post-infection followed a pattern that roughly matched the accumulation of progeny (**Fig. 1D**). The dsRNA staining was seen as clear puncta in SARS-CoV-2-infected cells, in a pattern suggestive of virus factories.

### Monocytes and T lymphocytes are the main targets of SARS-CoV-2 *in vitro* infection

To determine the susceptibility of circulating leukocytes to SARS-CoV-2, PBMCs from five healthy donors were infected (MOI=1), and analyzed the intracellular expression of SARS-CoV-2 antigens by flow cytometry. After 24 hpi, SARS-CoV-2 was detected in all immunophenotyped cells (**Fig 2A**). Monocytes were the most susceptible cell type, showing significant SARS-CoV-2 antigen staining (44.3%, p=0.039) (**Fig 2B**). In addition to monocytes, T CD4^+^ (14.2%, p=0.028), CD8^+^ (13.5%, p=0.019) and B lymphocytes (7.58%) were also susceptible to SARS-2 infection (**Fig 2C**). Staining for SARS-CoV-2 was significantly reduced in cells treated with NH_4_Cl, suggesting that acidification is important for in vitro infection of PBMCs.

### Infection of T lymphocytes leads to cell death by apoptosis

The COVID-19-related lymphocytopenia has been well described as a strong indicator of severe clinical outcomes in patients. Since we found both T CD4^+^ and CD8^+^ cells susceptible to SARS-2 infection *in vitro*, we investigated the presence of cell death in SARS-CoV-2-infected PBMCs from 5 healthy donors by the expression of translocated phosphatidylserine (PS) on the cell surface 24 hours post-infection by analizing its binding to annexin V (**Fig 3**). Despite the basal annexin V staining (CD4^+^ mean 6.24%, CD8^+^ mean 12.36%) seen in non-infected cells (**Fig 3A**), strong staining was observed both in live T CD4^+^ (70.88%, p=0.0001) and CD8^++^ lymphocytes (39.72%, p=0.0009) (**Fig 3B**). When cells were analyzed independently of Live/Dead staining, differences were still significant and even increased for CD8^+^ (59.64%, p=0.0001) (**Supplementary Fig 4**), indicating that a considerable percentage of Annexin V-positive CD8^+^ cells were already dead. No significant differences were observed in cell death between cells infected in the presence or absence of NH_4_Cl during infection. These results indicated that infection of human PBMCs by SARS-CoV-2 sharply increased the expression of apoptosis markers in T lymphocytes.

**Figure 4:**
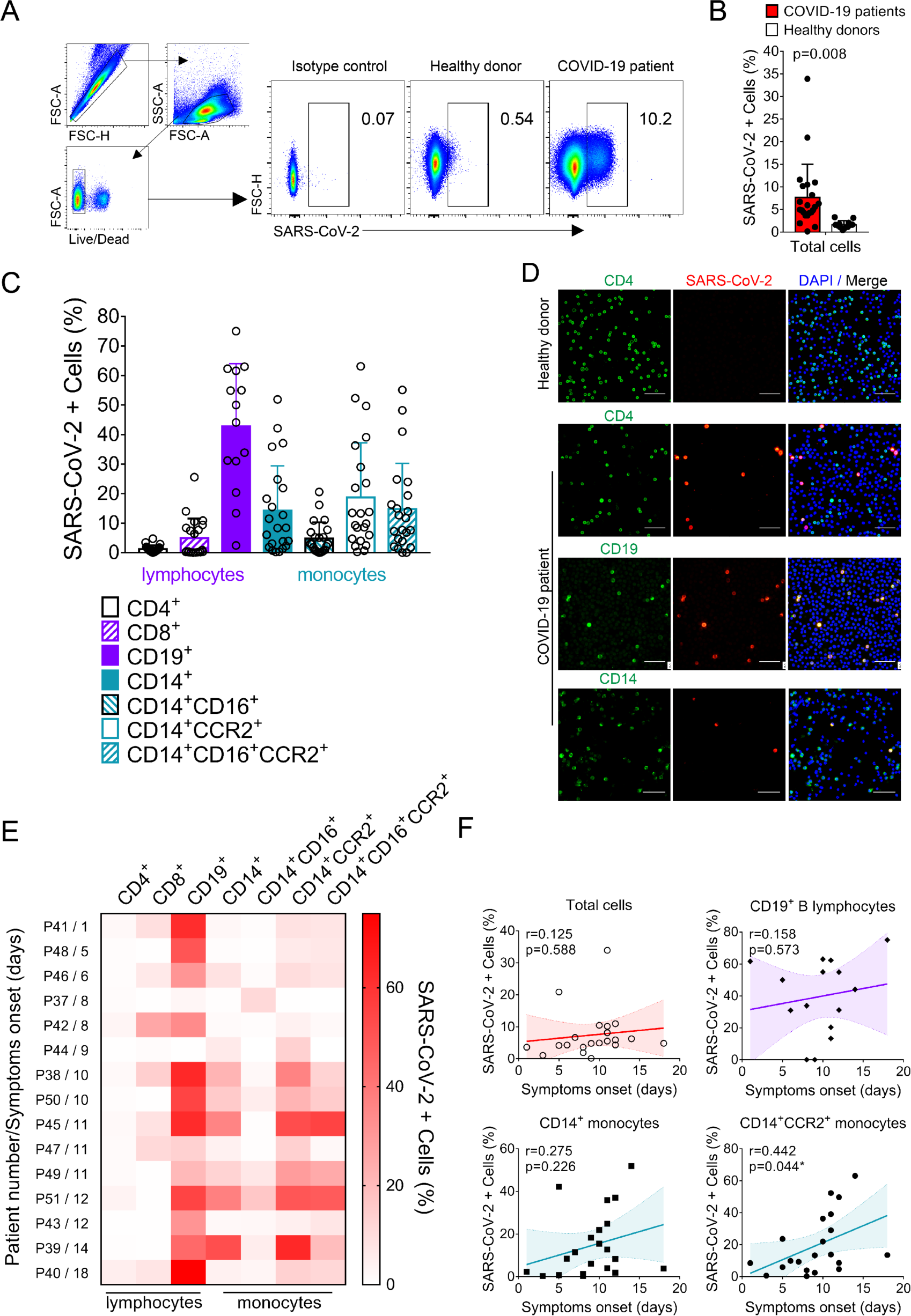
Detection of SARS-CoV-2 in PBMCs from hospitalized COVID-19 patients. (A) Representative flow cytometry plots indicating SARS-CoV-2 positivity of PBMCs from COVID-19 patients in comparison with isotype control and healthy donors. (B) Percentages of SARS-CoV-2-infected cells from COVID-19 patients (n=22) compared with background signal from healthy donors (n=12) cells. Results were compared by unpaired *t*-test (p=0.008). (C) Percentages of SARS-CoV-2-infected cells considering the different immunophenotypes, in COVID-19 patients (n=22). (D) Immunofluorescence of PBMCs from COVID-19 patients labeling for SARS-CoV-2 (red), nuclei (blue) and immunophenotypes CD4, CD19 or CD14 (green). Scale bars: 50 µM (E) Heat-map indicating SARS-CoV-2-positive cell frequencies for each immunophenotype, stratified by time from symptoms onset (Patient number/symptoms onset (days)). Data was plotted individually for each COVID-19 patient analyzed. (F) Correlation and linear regression analysis between time after symptoms onset and frequencies of SARS-CoV-2-positive cells. Both ‘p’ and ‘r’ values are indicated in the graphs. The best-fit line is displayed in all the graphs, while the light-color area represents the confidence interval. P values <0.05 were considered significant.

### Circulating immune cells from COVID-19 patients are infected by SARS-CoV-2

During April 7th to June 18^th^, we enrolled 22 COVID-19 patients that were admitted to the intensive care unit (ICU), presenting a moderate to severe disease. Clinical and demographic characteristics of enrolled patients are listed in Supplementary Table 1. Blood samples were collected at admission in the ICU. To check for SARS-CoV-2 infection in PBMCs from COVID patients, we analyzed PBMCs prepared from the whole blood of 22 patients and 11 healthy donors by flow cytometry (**Fig 4A**) with staining for SARS-CoV-2 antigens. Cells from COVID-19 patients showed significant expression of SARS-CoV-2 antigens (7.68%±1.56 p=0.008) in comparison with cells from healthy donors (**Fig 4B**). Interestingly, not all COVID-19 patients showed expressive staining for SARS-CoV-2, and rates of SARS-CoV-2-positive cells ranged from 0.16 to 33.9% (**Fig 4B**). Additionally, PMBCs from 15 COVID-19 patients were tested for the SARS-CoV-2 genome by real-time RT-PCR. Viral genome was detected in 8 out of 15 PBMC samples (53.3%), with mean viral load of 3.8×10^4^ copies per µg of RNA (Supplementary Table 2). Immunophenotyping of cells from COVID-19 patients indicated that the highest proportion of SARS-CoV-2-positive cells was found in B lymphocytes (42.73%±4.3). Although susceptible to *in vitro* infection, we were not able to find significant numbers of SARS-CoV-2 positive T cells in PBMCs from COVID-19 patients by flow cytometry. Similarly to what was observed by the *in vitro* experiments, monocytes (CD14^+^) from patients were found to be positive for SARS-CoV-2 in a high percentage (14.19%±15.26). Inflammatory monocytes (CD14^+^CCR2^+^ and CD14^+^CD16^+^CCR2^+^) were positive for SARS-CoV-2 antigen in rates significantly higher in comparison with healthy controls (18.73%±18.46 and 14.78%±15.5, respectively) (**Fig 4C**). To confirm the results obtained by flow cytometry, immunofluorescence was done for SARS-CoV-2 antigens in PBMCs isolated from COVID-19 patients. Some staining of SARS-CoV-2 with variable intensity was observed in CD19 and CD14 cells in PBMCs from COVID-19 patients, with no discernible fluorescent signal seen in PBMCs from healthy donors (**Fig 4D**). Despite what was observed by FC experiments, some IF staining was found in CD4 T lymphocytes, and after extensively screening, very few T CD8 cells were found to be positive for IF (**Supplementary Fig 5**).

**Figure 5.**
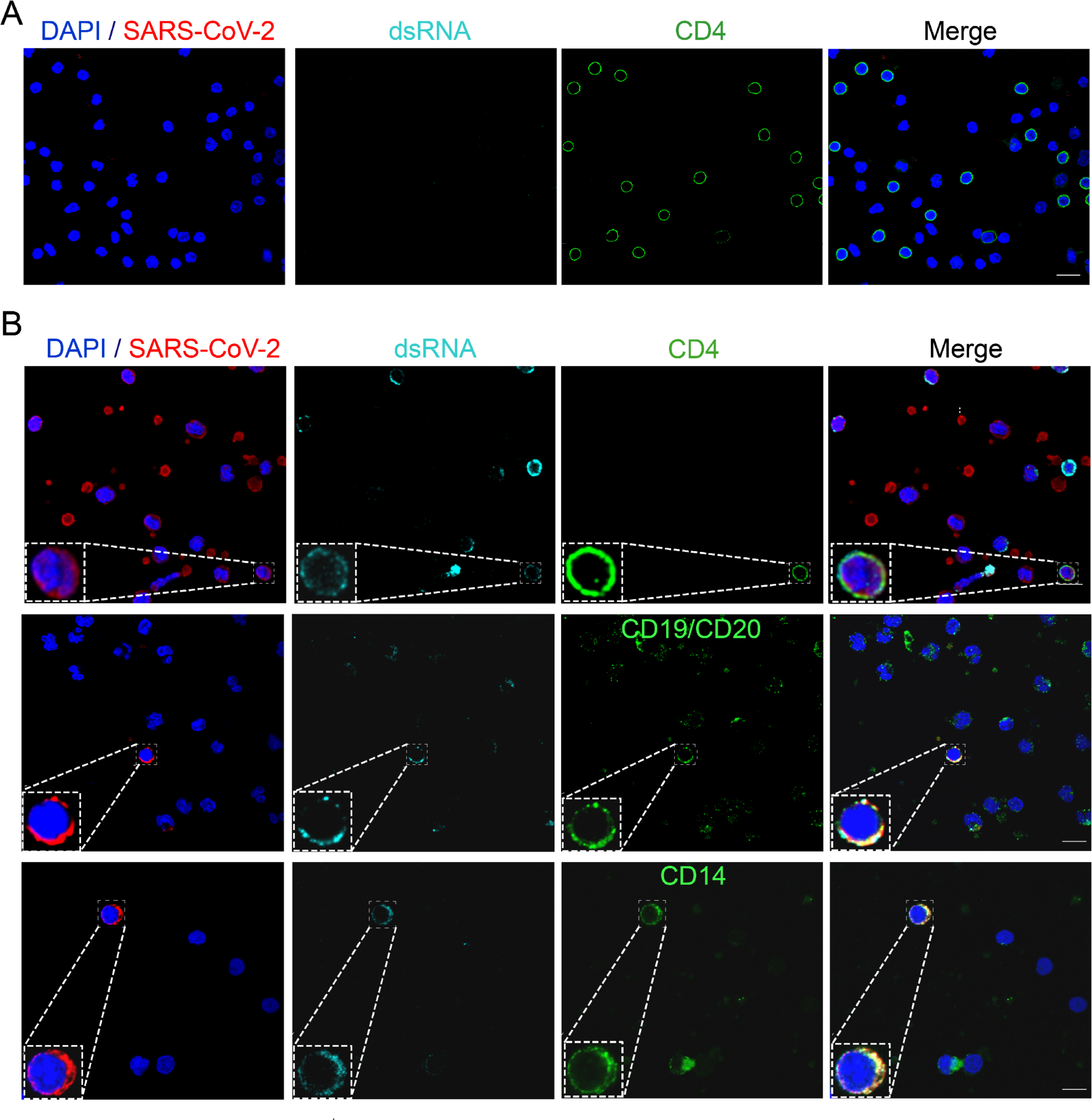
Peripheral blood cells naturally infected by SARS-CoV-2 from COVID-19 patients presents double-stranded RNA, a replication intermediate. PBMC from (A) healthy donors or (B) COVID-19 patients were isolated and put on coverslips pre-treated with poly-lysyne. Cells were fixed and stained for SARS-CoV-2 (red), immune phenotypes as CD4, CD19 or CD14 (green), dsRNA (cyan) and nuclei (blue). Immunofluorescence was examined using confocal microscopy. In the bottom left corner of each channel, an inset of the labelling phenotype is shown. Representative images for each immunophenotype, where at least two patients were analyzed. Magnification 63x. Scale bar 10 um.

Since the detection of SARS-CoV-2 in patients was found to be variable (Figure 4C), we selected 15 COVID cases to analyze individual differences in rates of SARS-CoV-2-positive cells. Patients were stratified based on the time of sample collection after symptoms onset, and SARS-CoV-2-positive cell frequencies were plotted on a heatmap for all cell immunophenotypes analyzed (**Fig 4E**). It became clear that rates of SARS-CoV-2-positive B lymphocytes were high throughout the entire dataset. In contrast, rates of SARS-CoV-2-positive monocytes were higher after following time progression after symptoms onset (**Fig 4E**). Frequencies of SARS-CoV-2-positive cells correlated positively with the length of time of COVID-19 progression after symptoms onset, especially for inflammatory CD14^+^CCR2^+^ monocytes (r=0.442 p=0.044) (**Fig 4F**).

To confirm whether SARS-CoV-2 was actively replicating in PBMCs from COVID-19 patients, we analyzed the presence of dsRNA in SARS-CoV-2-positive cells of different immunophenotypes by immunofluorescence and confocal microscopy. Remarkably, dsRNA staining was found in most SARS-CoV-2-positive cell subsets, CD4^+^ T lymphocytes, B lymphocytes, and monocytes (**Fig 5**). Altogether, these data confirm that SARS-CoV-2 infects circulating white blood cells from COVID-19 patients, and the frequencies of SARS-CoV-2-positive monocytes in the peripheral blood increase with time of onset of symptoms.

### Infected inflammatory monocytes are detected post mortem in lung tissues from COVID-19 patients

The respiratory tract is the classical entry route of coronaviruses in mammalian hosts. Therefore, we checked if the same infected cell immunophenotypes found in PBMCs could also be found by immunohistochemistry in the lungs of COVID-19 patients obtained *post mortem*. Post mortem lung specimens from COVID-19 patients revealed abundant staining for SARS-CoV-2, especially throughout the entire bronchovascular axes and alveolar-capillary barriers. Control lung specimens showed no staining (**Supplementary Fig 6**). Upon staining for SARS-CoV-2, slides were scanned, the staining was erased, and re-stained sequentially for the surface antigens CD4, CD20, and CD14. The serial immunolabelling indicated that CD4^+^ T lymphocytes, B lymphocytes, and monocytes express SARS-CoV-2 antigens (**Fig 6**) in the lungs of COVID-19 cases. Additionally, due to its well-known role in lung tissue damage in COVID-19, IL-6-positive cells were also searched for and, interestingly, several CD14^+^ monocytes expressing IL-6 were also positive for SARS-CoV-2 (**Fig 6C-E**), indicating that inflammatory monocytes in lungs of COVID-19 patients can also be infected with SARS-CoV-2.

**Figure 6.**
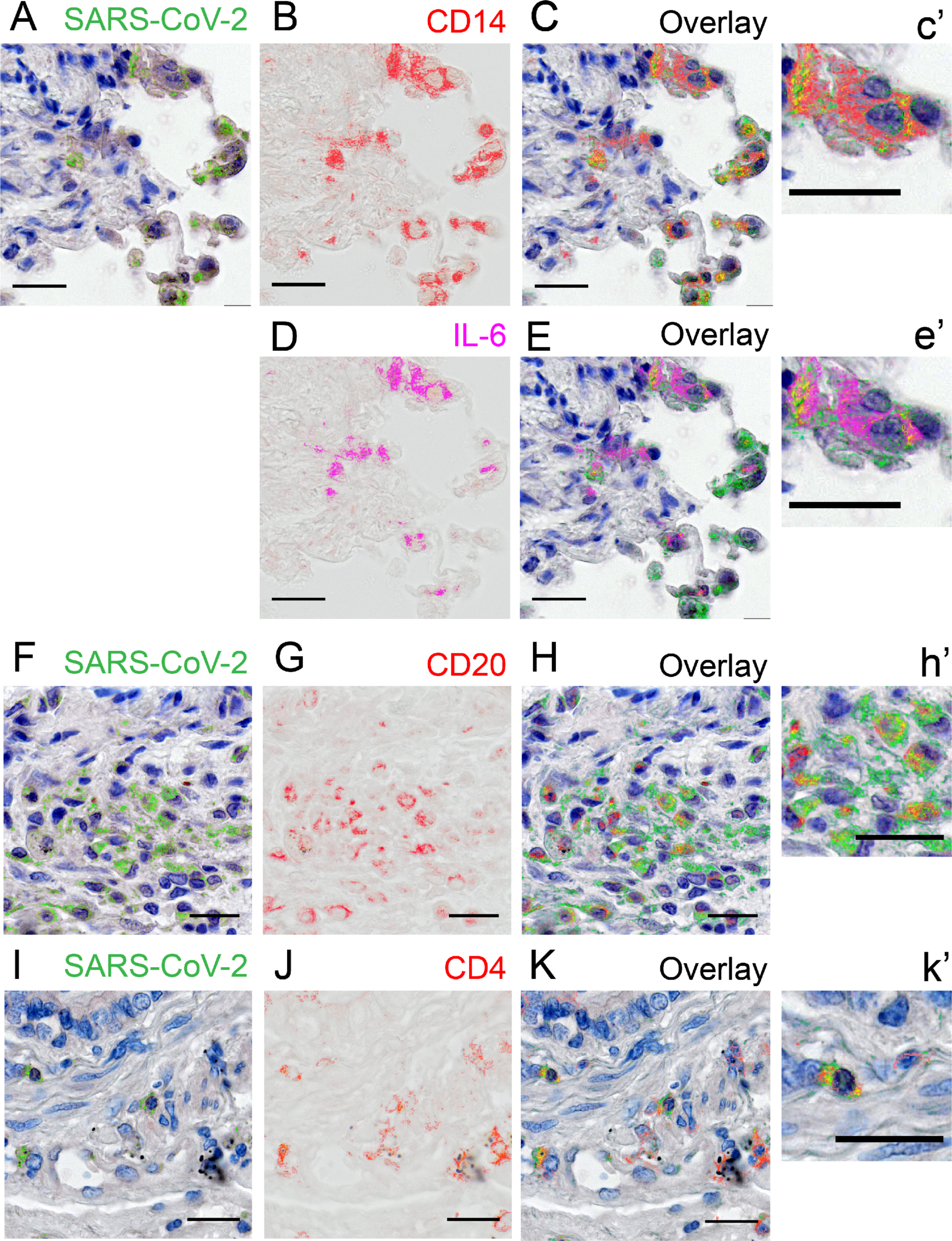
SARS-CoV-2 is detected in diverse immune cell types in COVID-19 lungs. (A, F and I) SARS-CoV-2 staining pseudocolored in green with hematoxylin counterstaining. (B, G, J) Staining for the immunophenotypes CD14, CD20 and CD4, respectively, pseudocolored in red. (D): Staining for IL-6, pseudocolored in magenta. (C, E, H and K) Overlaid layers from the previous sequential rounds of staining, with superimposed staining indicated in yellow. (c’, e’, h’ and k’) Insets from the respective previous panels. Scale bars: 50 µM.

## Discussion

It has been well accepted that several SARS-CoV-2 strategies to escape innate immune sensing, coupled with dysregulation of immune responses in early phases of infection, drive a cytokine storm that is a hallmark of severe COVID-19 [15–17]. Importantly, lymphopenia has also been recognized as a feature of severe infection by SARS-CoV-2. Postmortem examination of spleens and lymph nodes showed the presence of SARS-CoV-2 in those organs, infecting ACE2-expressing macrophages and causing important tissue damage [10].

SARS-CoV-2 detection in tissues far from the entry sites in the respiratory tract, without exuberant viremia, suggests that SARS-CoV-2 may reach target organs by alternative ways. One possibility could be the infection of leukocytes that could serve as “Trojan horses” transporting the virus to secondary infection sites. In that regard, we have recently reported that SARS-CoV-2 infects neutrophils, which could also act as Trojan horses carrying SARS-CoV-2 to neutrophil infiltrated tissues [18]. However, until now, it has been unclear whether SARS-CoV-2 infects PBMCs in vivo, thus creating a possibility of them being Trojan horses of viral dissemination. To address this question, we first infected PBMCs from healthy donors *in vitro* with SARS-CoV-2, as a preliminary way to check for their susceptibility and permissiveness to the virus. Virus production in PBMCs peaked at 12 hpi, reaching titers 100-fold the initial input, with steady decay thereafter until 48 hpi. The presence of dsRNA in SARS-CoV-2 infected PBMCs in the first few hours after infection provides further evidence that the virus replicates, yet modestly, in PBMCs in vitro. These results are in keeping with reports of SARS-CoV infection of human PBMCs [19]. Moreover, the treatment of PBMCs with ammonium chloride, which elevates the pH and prevents organelle acidification, significantly reduced but did not abrogate SARS-CoV-2 replication, consistent with an alternative acidification-independent pathway.

The immunophenotyping of PBMCs infected in vitro with SARS-CoV-2 revealed that CD14^+^, CD4^+^, CD8^+^ and CD19^+^ cells were susceptible. Primary human monocytes have been reported as susceptible to MERS-CoV and, more recently, to SARS-CoV-2 [20, 21]. In contrast, another recent study did not report PBMC infection by SARS-CoV-2 in vitro, possibly due to the low MOI used [22], coupled with the reported reduced expression of ACE2 by lymphocytes [14]. Despite that, we found that blockade of ACE2 partially reduced SARS-CoV-2 titers in supernatants of infected PBMCs, suggesting that among PBMCs there are ACE2-expressing cell types that contribute to the total virus progeny production. However, ACE2 blockade does not eliminate virus production, what strongly suggests the existence of ACE2-independent mechanisms of infection in lymphohematopoietic cells.

A recent report indicated that SARS-CoV-2 spike protein can interact with surface CD147, which could be an alternative virus receptor, in a way similar to what was observed for SARS-CoV [7, 23]. The transmembrane glycoprotein CD147, also known as Basignin, is expressed in some subsets of T lymphocytes [24], and thus could play a role in SARS-CoV-2 entry in these cells as well.

An intense T cell depletion in peripheral blood is seen in up to 85% of severe COVID-19 patients [2, 25]. Furthermore, T cells from COVID-19 patients show considerable levels of exhaustion markers [3, 4], and transcriptome analysis of their BBMCs indicated upregulation of genes involved in apoptosis and p53-signalling pathways [5]. These data suggests that SARS-CoV-2 infection could induce cell death by apoptosis in PBMCs, what could also happen in inflamed secondarily infected organs of COVID-19 patients. Of note, lymphopenia was also described in Middle East Respiratory Syndrome (MERS) patients, in whom MERS-CoV can directly infect human primary T lymphocytes and induce T-cell apoptosis through extrinsic and intrinsic pathways [20].

Annexin V staining showed that SARS-CoV-2 infection of PBMCs caused increased translocation of phosphatidylserine (PS) to the cell surface of both CD4^+^ and CD8^+^ T lymphocytes. The translocation of PS and subsequent scrambling of lipid membrane asymmetry is indicative of late-stage apoptosis [26]. Importantly, in the presence of NH_4_Cl, SARS-CoV-2 infection significantly increased annexin V labelling, suggesting that even at reduced levels of replication, SARS-CoV-2 can trigger apoptosis in lymphocytes. Taken together, the data indicate that SARS-CoV-2 infection of lymphocytes causes cell death, which may concur to the observed lymphopenia. The association of lymphopenia with poor prognosis may be related to the death of specific T-cell subsets, which may result in loss of immune response regulatory components, and drive a cytokine storm that can crosstalk with neutrophil NETosis [27]. Also, it can be related to increased IL-6 and Fas-FasL interactions [10], resulting in severe lymphoid tissue alterations [28].

In addition to *in vitro* infection, SARS-CoV-2 was also detected in PBMCs from COVID-19 patients, more prominently in B lymphocytes and subpopulations of monocytes. The predominance of B lymphocytes as target cells of SARS-CoV-2 infection *in vivo*, in contrast to what was seen in PBMCs infected *in vitro*, suggests that the susceptibility of different lymphocyte subsets in natural SARS-CoV-2 infection may depend on ACE2-independent alternative virus entry mechanisms. These findings corroborate previous observations that SARS-CoV enters B lymphocytes and monocyte-derived cells via a FcγRII-dependent pathway, which is facilitated by the presence of antibodies [29, 30]. The present results were obtained based on one-time sampling of patients who were enrolled at different times of COVID-19 evolution, what may explain the heterogeneity in rates of SARS-CoV-2-positive cells of different immunophenotypes observed among them. Accordingly, SARS-CoV-2 RNA was not detected in PBMCs from all, but in 53% of patients, indicating that SARS-CoV-2 infection in PBMCs may be variable, depending on host factors still unidentified, or present only in later phases of COVID-19, as suggested by the positive correlation between time from symptoms onset and frequency of SARS-CoV-2 positive cells in PBMCs. A possible explanation for an increase in SARS-CoV-2 susceptible cells over time could be an increase in ACE2 expression, triggered by type I IFN [31]. In this context, it is noteworthy that the replication of SARS-CoV in PBMCs was not sustained for long periods [19, 32].

To the best of our knowledge, this is the first report of circulating lymphoid cells positive for SARS-CoV-2, and the presence of dsRNA indicates that these cells are targets of virus replication. This may considerably impact the cells immune competence during COVID-19 and may help cell-associated SARS-CoV-2 spread to secondary infection sites.

SARS-CoV-2 recruits important inflammatory infiltrate in the lungs, containing diverse immune cell types that bear close contact with SARS-CoV-2-infected lung cells, such as pneumocytes and alveolar macrophages [33]. In the present study, we found CD4^+^ T and B lymphocytes and, importantly, also IL-6-expressing inflammatory monocytes positive for SARS-CoV-2 infiltrating the lung tissue from fatal cases of COVID-19. Further studies will be required to clarify whether SARS-CoV-2-positive lympho-mononuclear cells become infected in the lung or enter the affected tissue from the bloodstream already containing the virus. Regardless of where the immune cells become infected by SARS-CoV-2, their presence in the peripheral blood can impact directly on virus dissemination, delivering the infectious virus to secondary sites of infection.

Inflammatory monocytes play a significant role in the immunopathology of COVID-19 [15, 16] and ICU patients have high levels of circulating CD14+CD16+ inflammatory monocytes, which correlates with unfavorable outcomes [34, 35]. Increased expression of CCR2 and other inflammatory markers by monocytes leads to the infiltration of tissues with high expression of the correspondent CCL2 chemokine [15]. Interestingly, inflammatory monocytes with the same profile were found abundantly in bronchoalveolar lavage fluids from patients with severe COVID-19 [36]. Based on that, our data suggest that CD14+CCR2+ and CD14+CD16+CCR2+ infected monocytes could act as Trojan horses and traffic viruses to secondary sites of infection, where SARS-CoV-2 causes severe tissue damage. Additional lymphoid cell recruitment to damaged tissues may further contribute to lymphopenia [9].

Overall, the infection of lymphomononuclear cells by SARS-CoV-2 in peripheral blood from patients with COVID-19 has important consequences for pathogenesis of this multifaceted disease, including possible compromises of immune cell functions, and helping the virus to reach immune-privileged secondary sites of infection.

## Acknowledgements

This study was supported by the Brazilian National Research Council (CNPq grant numbers 310100/2017-8, 403201/2020-9 and INCTC 465539/2014-9) and the Sao Paulo State Research Foundation (FAPESP grant numbers 2013/16349-2 and 2014/02438-6). Dr. Pontelli was funded by CNPq grant number 380849/2020-8. The authors also thank Dimensions Sciences, a Non-Profit Organization that granted Dr. Castro a research scholarship while this study was conducted.

## Figure legends

**Figure S1.**
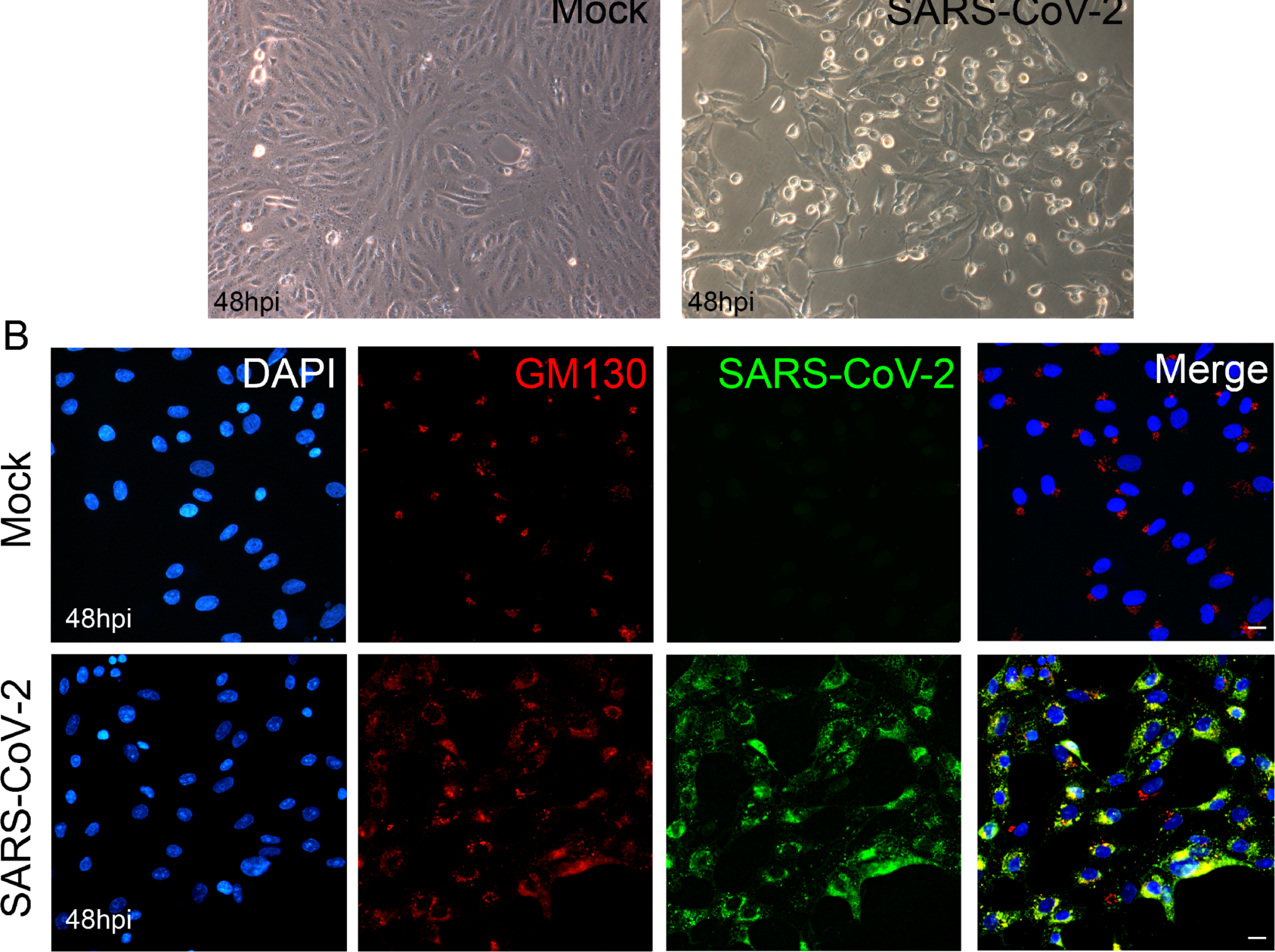
Validation of SARS-CoV-2 detection with human convalescent serum. Vero cells were infected with SARS-CoV-2 (MOI=1) or mock infected and incubated for 48h. (A) Phase-contrast microscopy of uninfected (left panel) and SARS-CoV-2-infected Vero cell monolayer showing cytopathic effect. Magnification 400×. (B) Immunofluorescence of Vero cells infected with SARS-CoV-2 or mock-infected at 48hpi, when cells were fixed and stained for GM130 (red), virus (green) and nuclei (DAPI). Scale bar 10 µm.

**Figure S2.**
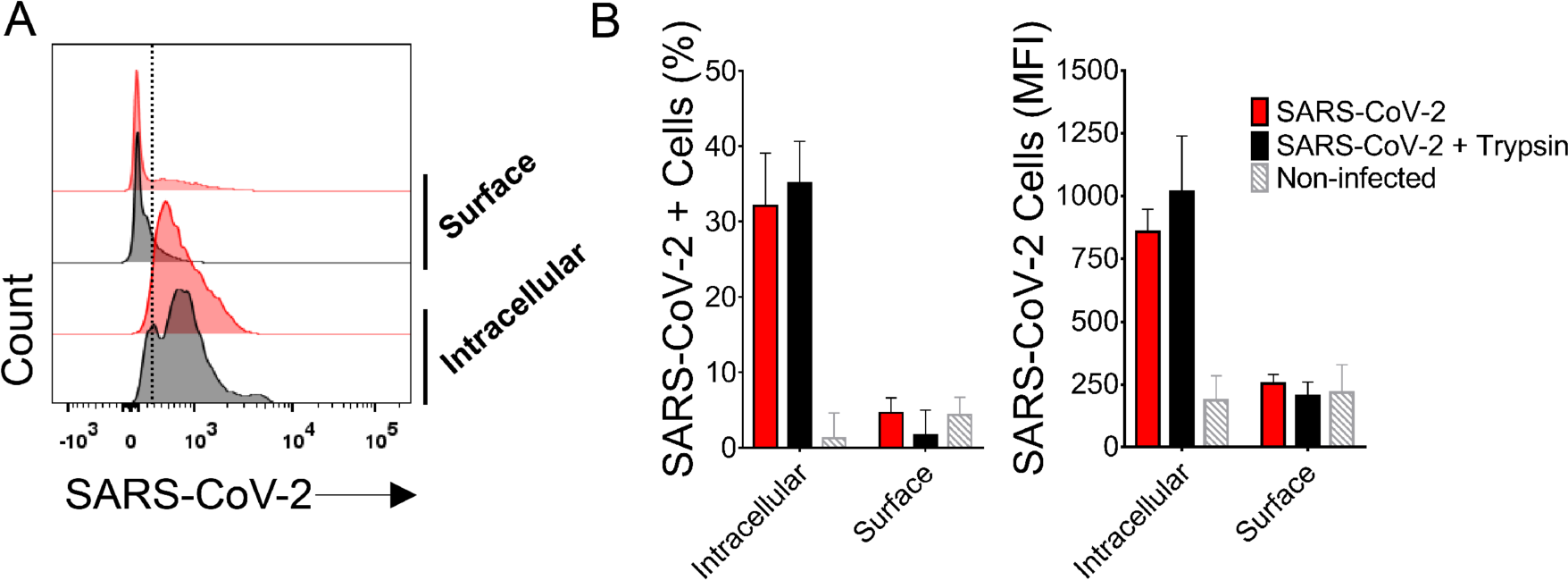
Flow cytometry (FC) of SARS-CoV-2-infected PBMCs from healthy with labeling for SARS-CoV-2. PMBCs from healthy donors infected in vitro (MOI=1) were analyzed by FC using mouse polyclonal anti-SARS-CoV-2 with and without cell permeabilization. Treatment with trypsin to remove surface-bound viral particles was used as an additional control. (A) Representative histograms of surface and intracellular staining for SARS-CoV-2, with SARS-CoV-2-infected cells in red and trypsin-treated infected cells in black. (B) Comparison of intracellular and surface staining of infected cells treated or not with trypsin, and non-infected cells in percentages on the left and MFI on the right.

**Figure S3.**
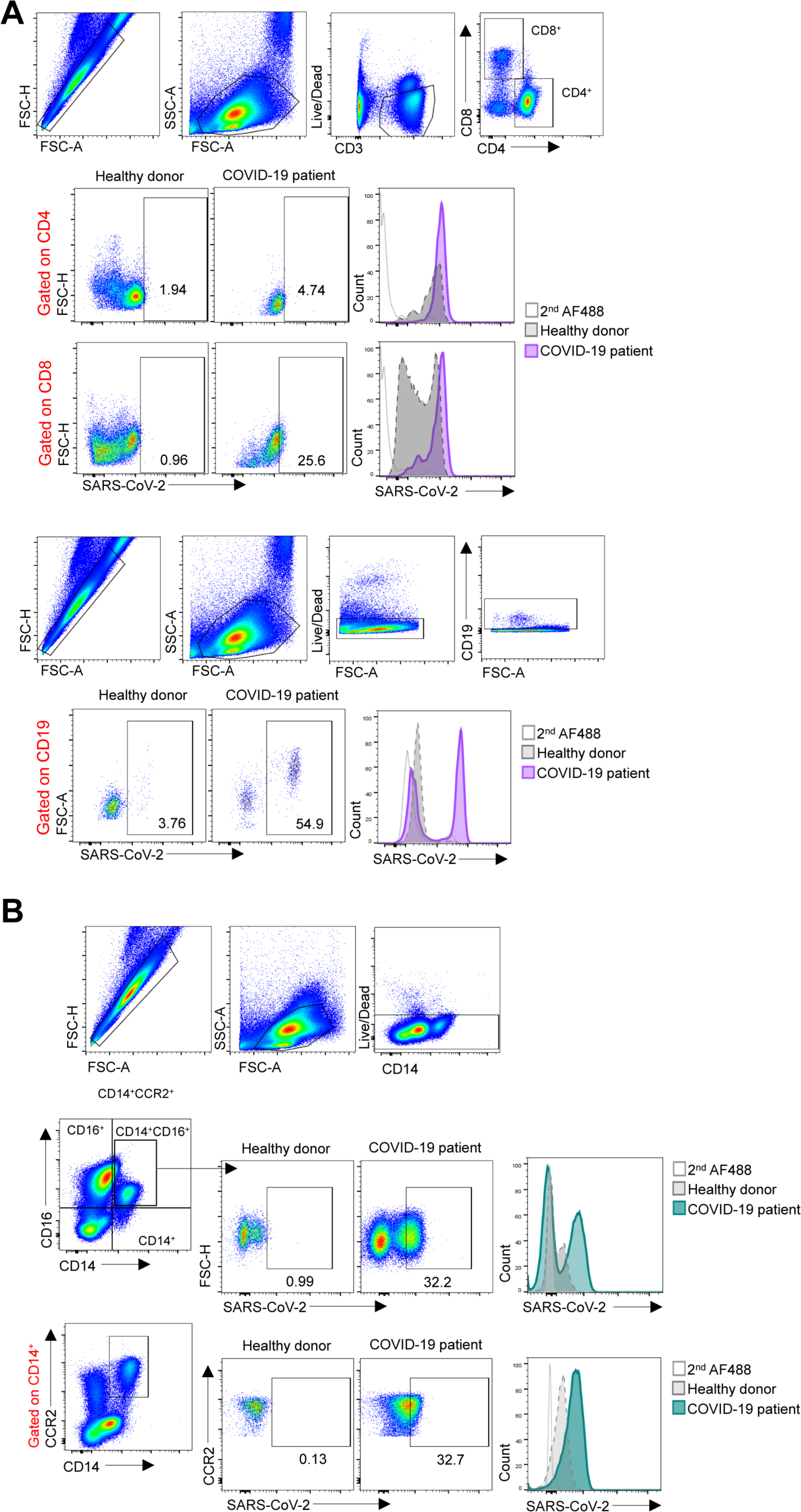
Gating strategies used for immunophenotyping of SARS-CoV-2-infected cells. (A) Cells were initially gated to exclude doublets and to exclude dead cells, using Live/Dead APC/H7 and CD3 staining. Next, detection of SARS-CoV-2 antigens in live T lymphocytes was defined based on the background secondary antibody signal (Alexa488) and signal obtained in healthy donors (flow plots and representative histograms). The same strategy was used for CD19^+^ B lymphocytes. (B) Live monocytes were initially gated as described for lymphocytes. Next, expression of CD14 and CD16 was used to define circulating monocyte subpopulations. Expression of CCR2 by CD14^+^ and CD14^+^CD16^+^ cells was used to define inflammatory monocytes. Among every defined subpopulation, expression of SARS-CoV-2 antigens was defined in comparison with secondary antibody background and healthy donors staining (flow plots and representative histograms).

**Figure S4.**
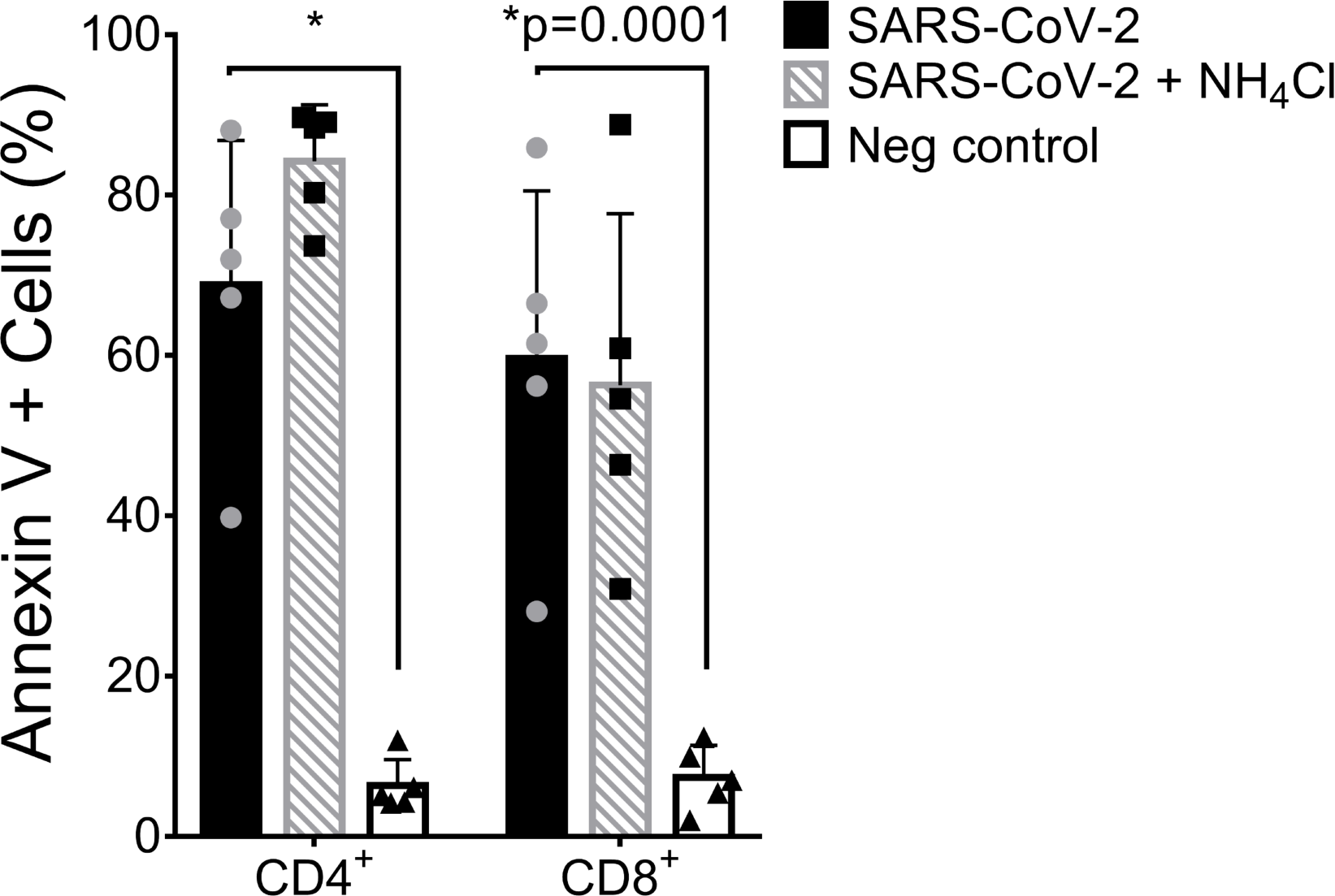
Percentage of lymphocytes expressing phosphatydilserine (PS) on the surface after in vitro infection with SARS-CoV-2. Cells were analyzed independently of Live/Dead APC/H7 staining. Mean ± s.d. is shown for all bar graphs. Significance was determined by one-way ANOVA and Bonferroni’s post-test was applied.

**Figure S5.**
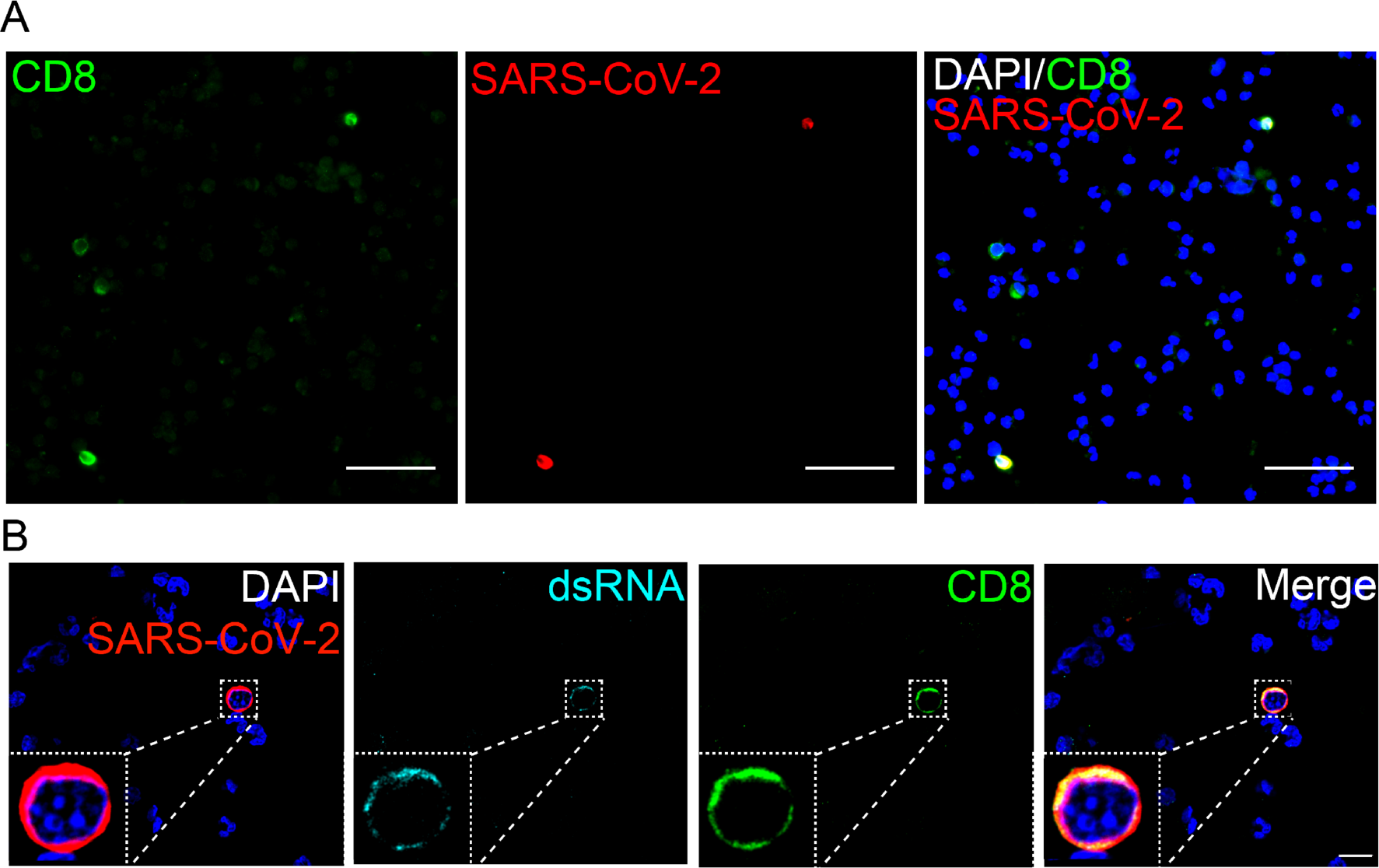
T CD8 lymphocytes are rarely detected with SARS-CoV-2. (A) PBMC from COVID-19 patients were put on coverslips pre-treated with polylysyne, fixed and stained for SARS-CoV-2 (red), CD8 (green) and nuclei (blue). Coverslips were analyzed in epifluorescence microscopy. Magnification 400x. Scale bar 20 um. (B) dsRNA detection in CD8 cells. PBMC was labeled as described in (a) and dsRNA (cyan) using an anti-J2 antibody. At the bottom left corner an inset is shown. Coverslips were analyzed in confocal microscopy. Magnification 63x. Scale bar 10 um.

**Figure S6.**
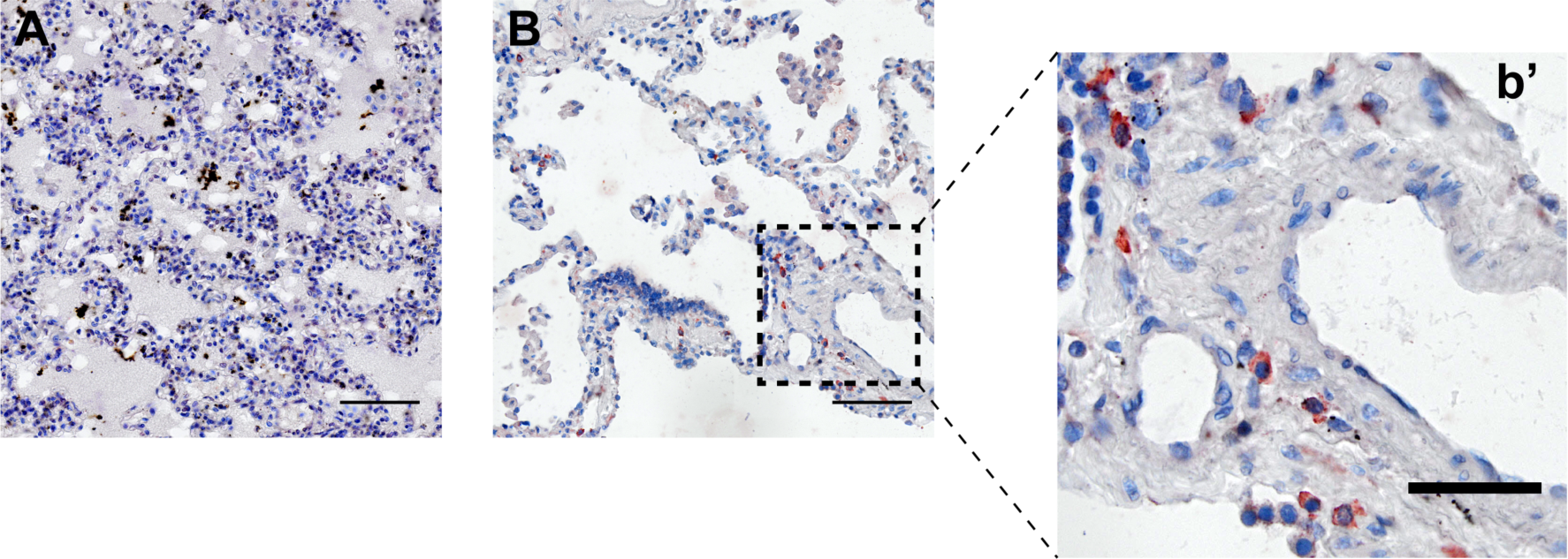
Immunohistochemistry for SARS-CoV-2 antigens in *post mortem* lungs from COVID-19. (A) *Post mortem* lung fragment from a hantavirus fatal case obtained in 2016, as a negative control for SARS-CoV-2 staining. (B) Staining for SARS-CoV-2 in lung from COVID-19 fatal case. (b’) Individual cells showing strong cytoplasmic staining for SARS-CoV-2 antigens in detail. Scale bars: 50 µM.

## Supplementary Tables

**Supplementary Table 1.** COVID-19 patient characteristics

| Demographics |  | % |
| --- | --- | --- |
| Number | 29 |  |
| Age (years) | 61.89±17.10 |  |
| Hospital day | 9.35±4.138 |  |
| Female | 13 | 44% |
| Comorbidities |  |  |
| Hypertension | 16 | 55% |
| Diabetes | 13 | 44% |
| Obesity | 13 | 44% |
| Lung disease | 4 | 14% |
| History of smoking | 9 | 31% |
| Heart disease | 7 | 24% |
| Kidney disease | 2 | 7% |
| History of stroke | 2 | 7% |
| Cancer | 4 | 14% |
| Autoimmune diseases | 2 | 7% |
| Immune deficiency | 2 | 7% |
| Laboratorial findings |  |  |
| CRP (mg/dL)* | 14.41±8.11 |  |
| D-Dimers (µg/mL)** | 2,244±1,698 |  |
| LDH (U/L)# | 749,4±490,5 |  |
| Ferritin (ng/mL)& | 1,985±2836 |  |
| Haemoglobin (g/dL) | 12.16±2.62 |  |
| Neutrophils (cell/mm<sup>3</sup>) | 7,521±4,952 |  |
| Lymphocytes (cell/mm<sup>3</sup>) | 1,666±1,286 |  |
| Platelets (count/mm<sup>3</sup>) | 231,009±136,433 |  |
| Image findings (n) | 29 | 100% |
| Medications |  |  |
| Antibiotics | 29 | 100% |
| Heparin | 29 | 100% |
| Antimalarial | 3 | 10% |
| Oseltamivir | 11 | 37% |
| Glucocorticoids | 16 | 55% |
| Respiratory status |  |  |
| Mechanical ventilation | 24 | 83% |
| Nasal-cannula oxygen | 27 | 93% |
| Room air | 0 |  |
| pO<sub>2</sub> | 70.36±45.64 |  |
| SatO<sub>2</sub> | 82.71±18.58 |  |
| Outcome |  |  |
| Deaths | 9 | 31% |

\*CRP: C-reactive protein (Normal value <0.5 mg/dL); \*\*D-dimers (NV <0.5 µg/mL); #LDH: lactic dehydrogenase (Normal range: 120-246 U/L); &Ferritin (NR: 10-291 ng/mL)

**Supplementary Table 2.** Viral loads of SARS-CoV-2 in PBMCs from COVID-19 patients tested by real-time RT -PCR

| Patient ID | Mean viral load |
| --- | --- |
|  | Genome copies/total RNA (µg) |
| P39 | 68560 |
| P42 | 22770 |
| P43 | 20782 |
| P45 | 36507 |
| P46 | 22140 |
| P47 | 94736 |
| P48 | 26455 |
| P50 | 16396 |
| **Mean±s.d.** | **38543±28114** |

